# Electrical stimulation in hippocampus and entorhinal cortex impairs spatial and temporal memory

**DOI:** 10.1101/215806

**Authors:** Abhinav Goyal, Jonathan Miller, Andrew J. Watrous, Sang Ah Lee, Tom Coffey, Michael R. Sperling, Ashwini Sharan, Gregory Worrell, Brent Berry, Bradley Lega, Barbara Jobst, Kathryn A. Davis, Cory Inman, Sameer A. Sheth, Paul A. Wanda, Youssef Ezzyat, Sandhitsu R. Das, Joel Stein, Richard Gorniak, Joshua Jacobs

**Affiliations:** Department of Biomedical Engineering, Columbia University, New York, NY 10027; Korea Advanced Institute of Science and Technology; Department of Biomedical Engineering, Drexel University, Philadephia, PA 19104; Thomas Jefferson University, Philadelphia, PA 19107; Mayo Clinic, Rochester, MN 55905; University of Minnesota, MN 55455; University of Texas, Southwestern, Dallas, TX 75390; Geisel School of Medicine at Dartmouth, Hanover, NH 03755; Hospital of the University of Pennsylvania, Philadelphia, PA 19104; Department of Neurology, University of Pennsylvania, Philadelphia, PA 19104; Department of Neurosurgery, Emory University, Atlanta, GA 30322; Department of Neurosurgery, Columbia University Medical Center, New York, NY 10032; Department of Psychology, University of Pennsylvania, Philadelphia, PA 19104

**Author notes:** Correspondence, 351 Engineering Terrace, Mail Code 8904, 1210 Amsterdam Avenue, New York, NY 10027.

## Abstract

The medial temporal lobe (MTL) is widely implicated in supporting episodic memory and navigation, but its precise functional role in organizing memory across time and space remains elusive. Here we examine the specific cognitive processes implemented by MTL structures (hippocampus and entorhinal cortex) to organize memory by using electrical brain stimulation, leveraging its ability to establish causal links between brain regions and features of behavior. We studied neurosurgical patients of both sexes who performed spatial-navigation and verbal-episodic memory tasks while brain stimulation was applied in various regions during learning. During the verbal memory task, stimulation in the MTL disrupted the temporal organization of encoded memories such that items learned with stimulation tended to be recalled in a more randomized order. During the spatial task, MTL stimulation impaired subjects’ abilities to remember items located far away from boundaries. These stimulation effects were specific to the MTL. Our findings thus provide the first causal demonstration in humans of the specific memory processes that are performed by the MTL to encode when and where events occurred.

**Significance Statement:** Numerous studies have implicated the medial temporal lobe (MTL) in encoding spatial and temporal memories, but they have not been able to causally demonstrate the nature of the cognitive processes by which this occurs in real-time. Electrical brain stimulation is able to demonstrate causal links between a brain region and a given function with high temporal precision. By examining behavior in a memory task as subjects received MTL stimulation, we provide the first causal evidence demonstrating the role of the MTL in organizing the spatial and temporal aspects of episodic memory.

## Introduction

The medial temporal lobe (MTL) plays a key role in encoding episodic memories for various types of spatial and temporal information (Eichenbaum, 2004; Cohen & Eichenbaum, 1993; Ekstrom et al., 2011). The importance of the MTL for memory is now well accepted, as researchers have reported concordant evidence from multiple methods, including observational studies of lesion patients (Scoville & Milner, 1957), experiments in rodents (O’Keefe & Dostrovsky, 1971), and, more recently, with data from noninvasive neuroimaging (Henson, 2005). However, although we know that the MTL is vital for episodic memory in general, we do not precisely understand the computational nature of the processes that the MTL employs to encode individual episodic memories in various contexts (Howard & Eichenbaum, 2015; Guderian et al., 2015; Maguire et al., 2015; Douglas et al., 2016).

Traditional methods for investigating the anatomical basis of human cognitive processes, such as lesion and neuroimaging approaches, have provided a plethora of information regarding the role of the MTL in memory (Bohbot et al., 2004; Copara et al., 2014; Suthana et al., 2009; Kolarik et al., 2017), but are unable to demonstrate causal links between a given brain region and a set of functions with high temporal precision. Localization of lesions is often uncontrolled, and the permanent nature of brain injury results in poor temporal resolution. Thus, one cannot always use lesion studies to perfectly identify the specific circumstances under which a given brain region is necessary for a given behavior (Rorden & Karnath, 2004). Analogously, neuroimaging studies are correlational, and are therefore unable to provide conclusive evidence about the necessity of a given brain region for a specific task in human subjects (Rorden & Karnath, 2004; Friston et al., 2002; Ramsey et al., 2010).

A different approach for probing brain function in humans is neuromodulation. Neuromodulation is promising because it can establish causal relationships between a brain region and particular behavioral functions (Knotkova & Rasche, 2014; S. H. Lee & Dan, 2012). Neuromodulation techniques such as optogenetics, transcranial magnetic stimulation (TMS), and electrical brain stimulation (EBS) allow researchers to transiently alter processing in a region and to determine the effects of this modulation on task performance (Suthana & Fried, 2014; Knotkova & Rasche, 2014). Showing that a cognitive process is transiently altered when a particular region is specifically stimulated demonstrates a causal link between the two (Suthana & Fried, 2014). Optogenetics is currently only implemented in animals, and TMS is only able to target neocortical structures (Klomjai et al., 2015). However, EBS is a viable research approach for certain neurosurgical patients, as it can target subcortical structures such as the hippocampus, making it a useful approach to directly map cognitive function in humans.

Previous work has shown that 50-Hz MTL stimulation impairs spatial and episodic memory overall (Jacobs et al., 2016). We move beyond the results of that work here by using an improved analytic approach on this dataset to identify the specific features of memory that were perturbed by stimulation. To foreshadow our results, we find that in the verbal domain, MTL stimulation disrupted the temporal ordering of episodic memory and impaired the recall of items from early list positions. Analogously, we found that stimulation specifically disrupted spatial memories for objects located far from boundaries, which we hypothesized were encoded with MTL-dependent representations, potentially based on grid or place cells. By performing quantitative, item-level analyses of behavioral data collected during brain stimulation, our findings provide the first causal and temporally precise demonstration of the specific cognitive processes that the MTL utilizes to organize episodic memories across time and space.

## Methods

We analyzed data from 49 epilepsy patients (32 female and 17 male) who had surgically implanted electrodes for localization of seizure foci as part of their evaluation for epilepsy surgery. Patients performed verbal-episodic and spatial memory tasks that we adapted such that direct brain stimulation was applied during some learning trials.

In each session, a selected electrode pair was connected to an electrical stimulator (Grass Technologies or Blackrock Microsystems). Stimulators were configured to deliver bipolar stimulation current between a pair of electrodes, each with a surface area of 0.059 cm^2^. MTL structures were stimulated using depth electrodes separated by either 5 or 10 mm, while strip and grid electrodes targeted other neocortical structures. Stimulation was applied at 50 Hz with a balanced biphasic stimulation pulse of 300 µs per phase for 1 second. Under clinical supervision, we began each session by manually testing a range of currents for each site, beginning at 0.5 mA and slowly increasing to a maximum of 1.5 mA (depth electrodes) or 3.0 mA (surface electrodes). The final stimulation current that was used for the cognitive experiments was the maximum current for each site that could be applied without inducing patient symptoms, epileptiform afterdischarges, or seizures.

We determined the anatomical location of each implanted electrode by examining an MRI scan, which provided a high resolution image of the hippocampus and MTL (0.5 mm × 0.5 mm × 2 mm resolution). Depth electrodes in the MTL were localized via a semi-automatic process. First, MTL subregions were labeled within the MRI using a multi-atlas based segmentation technique (H. Wang et al., 2013; Yushkevich et al., 2015). A neuroradiologist then identified each electrode contact on a post-implant CT scan, which was then co-registered with the MRI (Avants et al., 2008), and an anatomic label for each contact was automatically generated. Finally, the neuroradiologist visually confirmed the accuracy of the labeled location of the implant (Jacobs et al., 2016; Suthana et al., 2012). We designated a stimulation site as targeting a particular region if at least one electrode of the bipolar pair was in the region. Note that unlike our previous study, which analyzed stimulation effects at the level of individual sessions (Jacobs et al., 2016), here we analyzed stimulation-related changes to memory performance at the subject level, which is a slightly more conservative approach but nonetheless provided convergent results.

Each subject provided informed, written consent prior to participation. Our multisite study was approved by local institutional review boards (IRBs), the IRB of the University of Pennsylvania (data coordination site), and the Human Research Protection Official (HRPO) at the Space and Naval Warfare Systems Center Pacific (SPAWAR/SSC). The raw data are publically available (http://memory.psych.upenn.edu/) and were analyzed previously (Jacobs et al., 2016) but the results presented here are novel.

### Verbal Task

39 patients (23 with MTL stimulation) performed the free recall task (Sederberg et al., 2003) while stimulation was applied during the learning of some items. Figure 1A presents a timeline of this task. During each trial, subjects were asked to memorize a list of 12 words sequentially presented as text on the computer screen. Each word was presented for 1600 ms, followed by a blank screen for 750–1000 ms. Lists consisted of high-frequency nouns (http://www.memory.psych.upenn.edu/WordPools). After a 20-s math distractor task, subjects were given 30 s to verbally recall the words in any order. We recorded the verbal word responses for later manual scoring.

**Figure 1:**
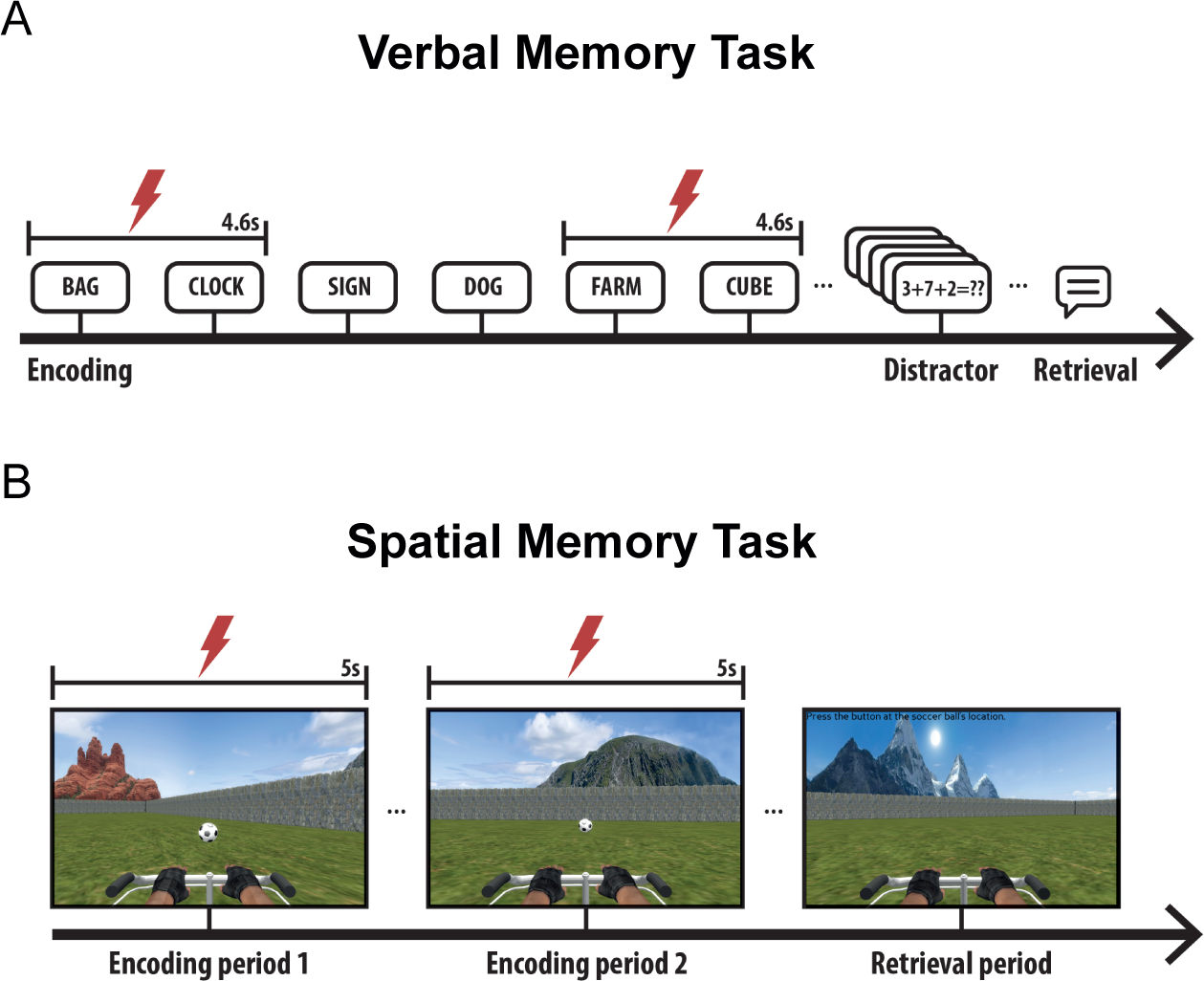
Overview of verbal and spatial memory tasks. **A.** Timeline of the verbal memory task. Half of the words on stimulated lists are encoded with the stimulator active. **B.** Overview of the spatial task. Half of the trials occurred with the stimulator active during both encoding periods. Taken from Jacobs et al. (2016) with permission.

Lists consisted of two types: stimulated lists, in which half of the words on the list were delivered simultaneously with electrical brain stimulation, and control lists, in which all twelve words on a list were presented without stimulation. Each session included 25 lists, consisting of 20 stimulated lists and 5 control lists in a random order. For each stimulated list, stimulation occurred in a blocked pattern: the stimulator was active during the presentation of a pair of consecutive words and then inactive for the following pair. Thus, in total, on each stimulated list the stimulator was active for half the total words. When the stimulator activated, it was timed to occur 200 ms prior to the presentation of the first word in each block, continuing for 4.6 s, until the disappearance of the second word. The onset of stimulation was balanced, such that a random half of the stimulation lists began with a non-stimulated block and the others began with a stimulated block.

### Data Analysis: Free Recall Task

Our analyses of data from the Free Recall task followed methods used in other studies (Kahana, 2012), adapted for examining the behavioral effects of stimulation. To examine how the effects of stimulation on memory varied over the timecourse of each stimulation cycle, we separately measured recall probability according to the position (“phase”) of an item within a stimulated or non-stimulated block. For this analysis, we averaged recall probability for each phase within stimulated and non-stimulated blocks and then normalized relative to matched positions on control lists.

For some analyses we separately measured memory performance as a function of list position. We defined the primacy period as the first two items on each list, consistent with previous free recall studies using similar list lengths (Fischler et al., 1970; Craik, 1970). This classification fit the data because, after drops in performance between the recall rates for the first two list positions of more than 0.06, memory performance was roughly comparable across positions 3–5 (changes in recall rates of less than 0.025).

To assess the effect of stimulation on erroneous recalls, we measured the rates of prior list intrusions (PLIs) (Darley & Murdock, 1971), which occurred when subjects incorrectly recalled an item from the previous list during the current list’s recall period. Many PLI probabilities were close to zero, therefore we assessed signifiance using nonparametric statistics.

In free recall, subjects exhibit a strong tendency to cluster recall sequences based on the temporal proximity of the items during study. We examined the effect of stimulation on this effect clustering by computing two measures of temporal clustering: temporal clustering factors (TCF) (Polyn et al., 2009b) and conditional response probabilities (CRPs) (Kahana, 1996). The TCF is a number that measures the mean tendency for recall transitions to occur between items presented at nearby list positions. A TCF of 1 indicates perfect temporal contiguity, with the subject only making transitions to temporally adjacent items, while a TCF of 0.5 indicates that the subject is making transitions randomly (Polyn et al., 2009a). The CRP is a curve that indicates the conditional probability of a particular item being recalled as a function of the difference in its position in the learned list relative to that of the previously recalled item. We performed all TCF and CRP comparisons with non-parametric tests. To rule out the possibility that differences in mean recall rates between stimulated and control lists led to apparent changes in TCFs, we used a downsampling procedure to artificially match recall rates between stimulated and non-stimulated lists.

We also examined the effects of stimulation on non-MTL regions. 5 subjects received stimulation in the frontal lobe, 2 in the cingulate, and 2 in the lateral temporal lobe. We termed these “neocortex” regions. To identify potential results caused by sample size differences between the MTL and neocortex datasets, we reperformed our analyses using a bootstrap procedure. In this procedure we chose a random subset of 9 MTL electrodes and then computed the effect of interest and confidence intervals across 1,000 iterations.

### Spatial Task

In the spatial memory task (Fig. 1B), 26 subjects received MTL stimulation while participating in a virtual navigation paradigm that is reminiscent of the Morris (1984) water maze. Eight of these subjects also participated in the verbal task with MTL stimulation. The environment was rectangular (1.8:1 aspect ratio), and was surrounded by a continuous boundary. There were four distal visual cues (landmarks), one centered on each side of the rectangle, to aid with orienting. Each trial started with two 5-s encoding periods, during which subjects were driven to an object from a random starting location. At the beginning of an encoding period, the object appeared and, over the course of 5 s, the subject was automatically driven directly towards it. The 5 second period consisted of three intervals: first, the subject was rotated towards the object (1 s), second, the subject was driven towards the object (3 s), and, finally, the subject paused while at the object location (1 s). After a 5-second delay with a blank screen, the same process was repeated from a different starting location. Alternating trials (24 out of the 48) were designated as stimulation trials, during which stimulation was applied throughout the 5 s of time while the object was visible to the subject for both encoding periods.

After both encoding periods for each item, there was a 5-s pause followed by the test period. The subject was placed in the environment at a random starting location with the object hidden and then asked to freely navigate using a joystick to the location where they thought the object was located. When they reached their chosen location, they pressed a button to record their response. They then received feedback on their performance via an overhead view of the environment showing the actual and reported object locations. Between stimulation and non-stimulation trials, starting location, starting orientation, and object location were all counterbalanced. This was achieved by creating a set of location triads for the stimulated conditions and transposing them across the environment’s diagonal for use in non-stimulation trials, ensuring that the geometric relationship between the start and object locations was matched in stimulation and non-stimulation trials.

### Data Analysis: Spatial Memory Task

Performance in the spatial task was measured by computing the distance between the reported and actual locations for each object. In the same manner as Jacobs et al. (2016), we normalized this euclidean distance metric into a memory score (MS) between the range of 0 and 1, where a value of 1 indicates a perfect response and a value of 0 indicates the worst possible response given the object’s location. This normalization took the form of ranking the subject’s actual response compared to all other possible responses. This normalization procedure removes potential bias in the results by accounting for the fact that the distribution of possible response error distances varies according to the object’s distance from the boundaries; it also corrects for the rectangular shape of the environment. Namely, objects near boundaries had a larger maximum possible error distance than objects in the interior. Subjects with an average MS of less than chance (0.5) were excluded from all analyses. We utilized Tukey’s honest significant difference (HSD) correction for multiple comparisons when performing post-hoc analyses of analysis of variance tests.

Previous studies have shown that boundaries play a crucial role in guiding navigational behavior (Chan et al., 2012; S. A. Lee et al., 2018; Hartley et al., 2004; S. A. Lee, 2017), so we chose to analyze the effect of object location on subject performance. To this end, we divided the environment into “boundary” and “interior” regions of equal area by creating an inner rectangle with an identical aspect ratio to the environment itself.

We hypothesized that subjects sometimes utilized view-based spatial memory strategies, which rely on facing the same direction during encoding and recall. Such strategies would be most effective where salient visual scenes were most prominent, which occurred when the subject was close to the environment’s boundaries. To test whether some subjects might have used such a strategy, we labeled the directions that subjects faced at the end of the learning trials and the test trials as the “headings” for that trial. The circular mean of the two learning headings was then subtracted from the test heading to compute the trial’s learning–test heading difference. All heading statistics were calculated with circular statistics (Berens, 2009).

We conducted a simulation to assess the possibility that our task’s design led to artificially decreased heading differences between learning and test for certain areas of the environment. In a single iteration of this procedure, we simulated 1,000 learning and test trials, with randomly generated start and end locations. For each simulated test trial, the simulated “subject” drove in a straight line from the start to the end location. Then, across these 1,000 trials, we computed the mean vector length (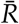) of the trial-wise learning–test heading differences. This entire procedure was then repeated 100 times to establish confidence intervals for 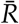.

## Results

We sought to determine how electrical stimulation of the hippocampus and entorhinal cortex influences the precise temporal and spatial organization of memory. To answer this question, we conducted new analyses of a previously published dataset (Jacobs et al., 2016) in which subjects performed spatial and episodic memory tasks with concurrent brain stimulation in the MTL. Going beyond the previous study, which reported that MTL stimulation reduced verbal and spatial memory performance overall, our new analyses show that MTL stimulation selectively disrupted the temporal organization of verbal memory and the ability to encode spatial locations in an environment without visual cues from boundaries.

### Verbal Episodic Memory

In the verbal memory task, subjects learned two types of word lists: stimulated lists and control lists. On stimulated lists, electrical stimulation was present for alternating blocks of two items at a time. Therefore, items on stimulated lists consisted of two categories: those delivered with stimulation (“stimulated items”) and those delivered without (“non-stimulated items on stimulated lists”). Control lists consisted entirely of items delivered without stimulation. Our data analyses separately examined recall rates across items from different categories. As reported in Jacobs et al. (2016), recall rates were lower for stimulated items relative to non-stimulated items (*t*_22_ = −2.29, *p* = 0.04, paired *t*-test), indicating that MTL stimulation impaired memory encoding. Going beyond this earlier work, we examined the time course of the effects of stimulation and whether stimulation affected memory for the order of learned items.

### Effect of Stimulation on Verbal Memory Encoding

First, we examined whether the memory impairment from stimulation lingered after stimulation ended. To do this, we measured recall performance for non-stimulated items on stimulated lists, as well as for items on control lists (Fig. 2A). We found significant differences across all three conditions (one-way ANOVA; *F*(1, 22) = 3.92, *p* = 0.020). Recall rates were lower for stimulated items compared to non-stimulated items (HSD-corrected post-hoc *t*_22_ = −2.29, *p* = 0.04, paired *t* test) and for non-stimulated items on stimulated lists relative to items on control lists (HSD-corrected *t*_22_ = −1.73, *p* = 0.07, paired *t* test). These results indicate that the memory impairment from stimulation persists after the stimulation interval, moderately impairing recall rates for items learned right after the stimulator was turned off.

**Figure 2:**
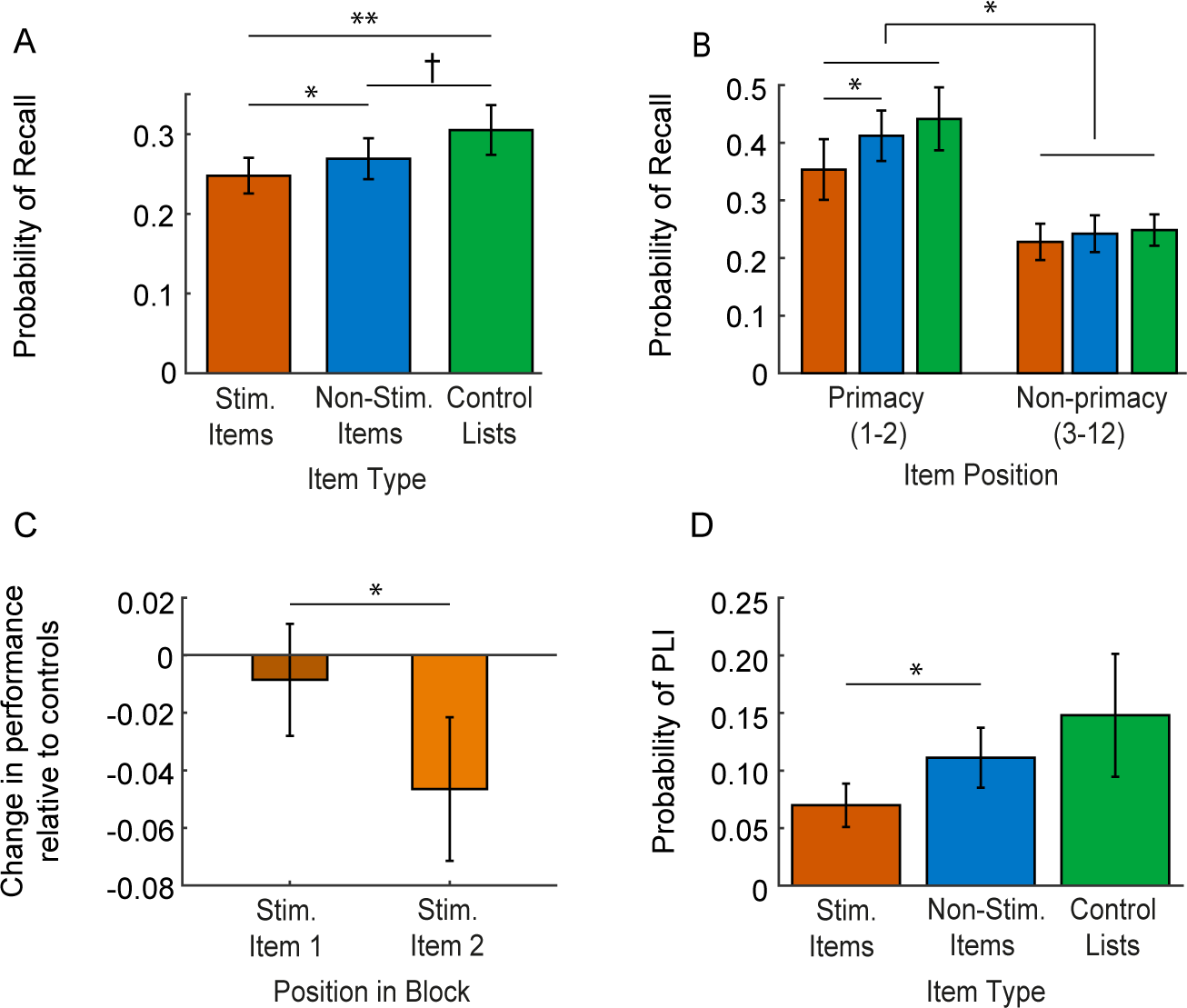
Effect of MTL stimulation on verbal memory encoding in free recall. **A.** Probability of item recall averaged across all serial positions, separately computed for stimulated items (red), non-stimulated items on stimulated lists (blue), and control lists (green), then averaged across subjects. **B.** Recall probabilities separately computed for primacy (items 1–2) and non-primacy items (items 3–12) in the same way as Panel **A. C.** Change in mean recall probability for stimulated lists versus control lists (position matched), computed relative to stimulation onset. **D.** Probabilities of incorrectly recalling items from different sources (labeled “item type”) while recalling items on a stimulated list. *: *p* ≤ 0.05, **:*p* < 0.01, †: *p* < 0.1. Error bars are SEM.

Electrophysiological studies have suggested that different neural patterns underlie the encoding of items from early and late positions within an individual list (Serruya et al., 2014; Sederberg et al., 2006; Manning et al., 2011). To compare the role of the MTL in encoding early versus late items, we measured the impact of stimulation across the course of each list. We computed recall probabilities for each stimulation condition separately for primacy (items 1 & 2; see *Methods*) and for non-primacy items (items 3–12; Fig. 2B). As expected (Murdock, 1962), overall recall probabilities for primacy items were higher than for non-primacy items. However, the effects of MTL stimulation varied over the course of the list. To assess this effect, we performed a two-way repeated-measures ANOVA with factors list position (primacy/non-primacy) and stimulation condition (stim., non-stim. item on stim. lists, and control lists). We found that MTL stimulation impaired the recall of items more for primacy than non-primacy positions (ANOVA interaction *F*(1, 44) = 2.78, *p* = 0.047). We confirmed that stimulation significantly impaired recall of primacy items (HSD-corrected post-hoc *t*_22_ = −1.95, *p* = 0.04, paired *t* test) and that this impairment was not present for non-primacy items (*p* > 0.8, *t* test).

As mentioned above, on stimulated lists the stimulator was enabled in a two-on-two-off cycle across items. To examine how memory performance varied according to the phase of the stimulation cycle, we compared the effect of stimulation on memory performance in these intervals relative to position-matched controls (see *Methods*). Memory performance was more strongly impaired for the second stimulated item compared to the first such item (Fig. 2C; *t*_22_ = −2.10, *p* = 0.042, paired *t* test), thus indicating that the impairment of memory from stimulation accumulates gradually or has a delayed onset.

In addition to comparing mean accuracy rates, an additional way to assess the effects of stimulation on memory is to investigate the types of errors that are made during recall (Darley & Murdock, 1971). To test whether stimulation influenced the types of recall errors that subjects made, we examined prior list intrusions (PLIs), defined as recalls of items from the previous list rather than the current one (Fig. 2D). We found that stimulated items on a previous list had a lower probability of being the source of a PLI compared to non-stimulated items on a previous list (*z* = −2.12, *p* = 0.034, *n* = 23, Signed Rank Test). This finding suggests that when an item is learned in the presence of MTL stimulation, it is less strongly maintained in memory.

### Effect of Stimulation on the Temporal Organization of Memory

We next examined whether MTL stimulation altered the temporal structure of memory. In a standard delayed free recall task without stimulation, subjects tend to recall items in the order that they were encoded (Howard & Kahana, 1999). We hypothesized that stimulation might disrupt this phenomenon. To examine this issue, we present the results of two separate analyses of the effect of stimulation on the temporal structure of episodic memory recalls. We begin by examining recall order at the beginning of each list, followed by a broader analysis of the overall temporal order of responses across the complete list (Polyn et al., 2009a).

To examine the effects of MTL stimulation on recall order, we computed the mean probability of recalling an item from each list position in each of the first four output positions (Fig. 3). On control lists, as expected, there was a tendency for items to be recalled in the order they were viewed. However, MTL stimulation disrupted this pattern, as we show by comparing recall rates at each position between control and stimulation lists (Figs. 3A–D). Subjects exhibited decreased probabilities for recalling each of the first four items in their proper orders (items 1, 3,& 4: 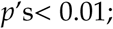 item 2: *p* = 0.078). Notably, when subjects recalled three or more items on a stimulated list, they most often recalled the third item first. These results indicate that stimulation hindered subjects from encoding temporal structure during learning.

**Figure 3:**
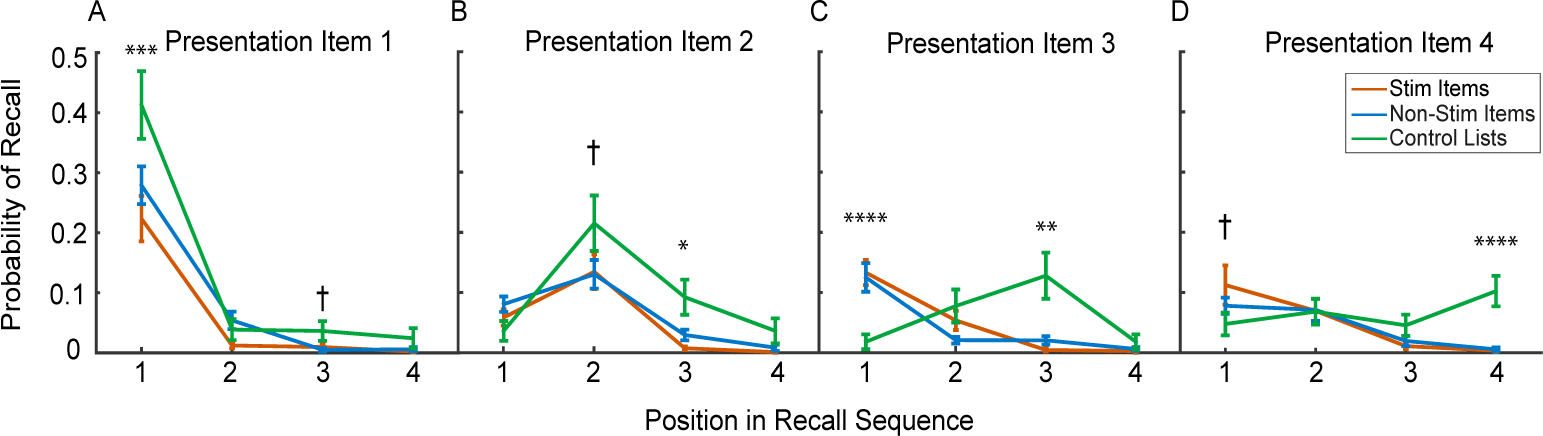
Effect of MTL stimulation on recall order for the beginning of the list. Plots show the probability of recall at different recall-output positions. Probabilities were averaged within each subject and then averaged across subjects. Plots from left to right show results separately for items that were presented at positions 1–4. *: *p* < 0.05, **: *p* < 0.01, ***: *p* < 10^−3^, ****: *p* < 10^−4^, †: *p* < 0.1

In the free recall task, item recalls tend to be temporally clustered, such that items consecutively recalled are more often learned from nearby list positions (Howard & Kahana, 2002). We examined the effect of stimulation on temporal clustering by computing two measures of list-wide temporal contiguity—the temporal clustering factor (TCF) (Polyn et al., 2009a) and the lag conditional recall probability (CRP) function (Kahana, 1996)—and testing if they changed with MTL stimulation (Fig. 4A). TCFs, which measure the correlation between item ordering during encoding and recall, were higher for control lists compared to conditions with stimulation (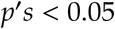, rank-sum tests). Despite the theoretical insensitivity of the TCF to recall counts (Polyn et al., 2009a), to rule out the possibility that the temporal factor was lowered by the diminished recall rates on stimulated lists, we reperformed this analysis after matching recall counts between control and stimulated lists with random subsampling. However, this analysis replicated the same pattern of results 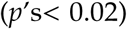, confirming our original interpretation that stimulation specifically impaired temporal clustering in addition to diminishing the mean recall rate.

**Figure 4:**
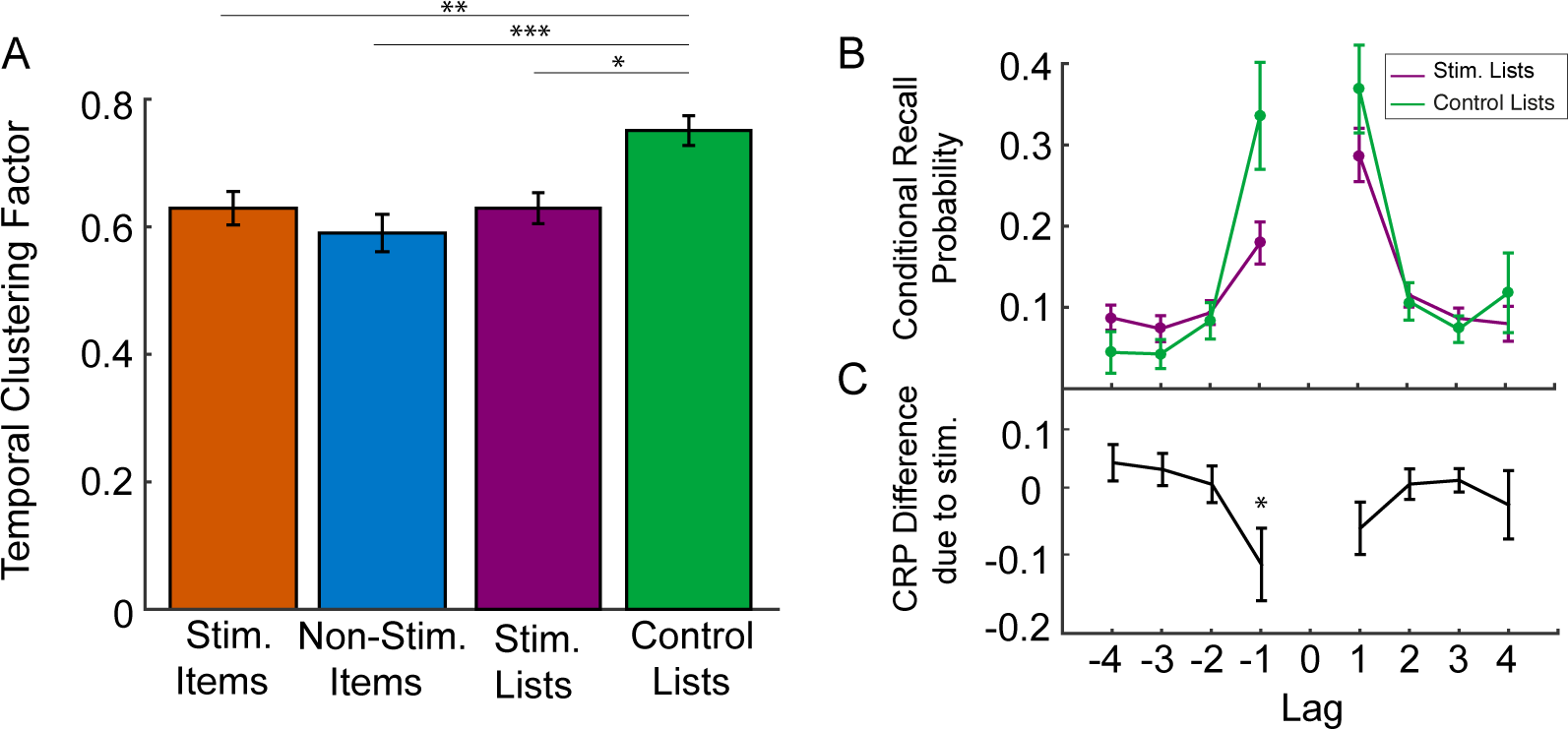
Analysis of the effect of MTL stimulation on temporal clustering of item recalls. **A.** Temporal Clustering Factors (TCF) for items from each stimulation condition, averaged across subjects. **B.** Conditional Recall Probability (CRP) plot for control and stimulated lists, averaged across subjects. **C.** Difference in recall probability from stimulation (Stim List control). *: *p* < 0.05,**: *p* < 0.01, ***: *p* < 10^−3^

To visualize the effect of MTL stimulation on the dynamics of memory, we computed the lag-CRP for each list condition. Overall, both stimulated and control lists show higher recall probabilities at short lags, as expected (Kahana, 1996). However, CRPs for stimulated lists were flatter than for control lists (Figure 4B). In particular, with stimulation there was a significant decrease in recall probability for item transitions at lag=−1 (*p* = 0.032, rank-sum test; Fig. 4C). These results support the notion that MTL stimulation disrupts the temporal organization of memory, by decreasing subjects’ tendencies both to recall items in their viewed order and to make temporally contiguous recalls.

An unexpected feature of our data was that the CRPs for control lists were rather symmetric, as opposed to showing a moderate asymmetry (Kahana, 1996). To explain this pattern, we separately examined CRPs for items from different list positions. The CRP for the first half of each list showed a normal forward asymmetry for both control and stimulated lists, and a significantly lowered recall probability at lag=−1 for stimulated compared to control lists (*p* < 0.05, rank sum test). In contrast, the CRP for the second half of the list was symmetric and showed no significant differences between stimulation and control conditions (*p* > 0.1). Thus, the symmetry of the aggregate CRP was caused by recalls in the second half of lists deviating from expected patterns.

### Stimulation outside the MTL

To compare whether the memory modulation patterns we observed were specific to stimulation in the MTL, we compared recall performance between sessions where stimulation was applied in MTL regions (hippocampus and entorhinal cortex) versus stimulation outside the MTL (neocortex). Recapitulating past results (Jacobs et al., 2016), neocortical stimulation did not significantly impair recall rates compared to control lists (Fig. 5A; *p* > 0.1, one-sample *t* test). This effect was significantly different compared to the effect of MTL stimulation (*t*_31_ = −1.76, *p* = 0.044, unpaired *t* test). Within the neocortex group, stimulation electrodes were placed in different subregions across subjects (see *Methods*). However, we did not observe significant differences in the effects of stimulation between these subregions (one-way ANOVA, *F*(2) = 0.73,*p* = 0.52). There was also no significant effect of neocortical stimulation on temporal clustering (Fig. 5B). Thus, at least with the type of stimulation we used, stimulation’s ability to disrupt the temporal organization of memory is specific to MTL sites and is not a brain-wide phenomenon. Finally, we note that these regional differences were not a result of differing sample sizes between MTL and neocortex, because we found the same pattern of results after reperforming this analysis with sample sizes matched using random subsampling (see *Methods*).

**Figure 5:**
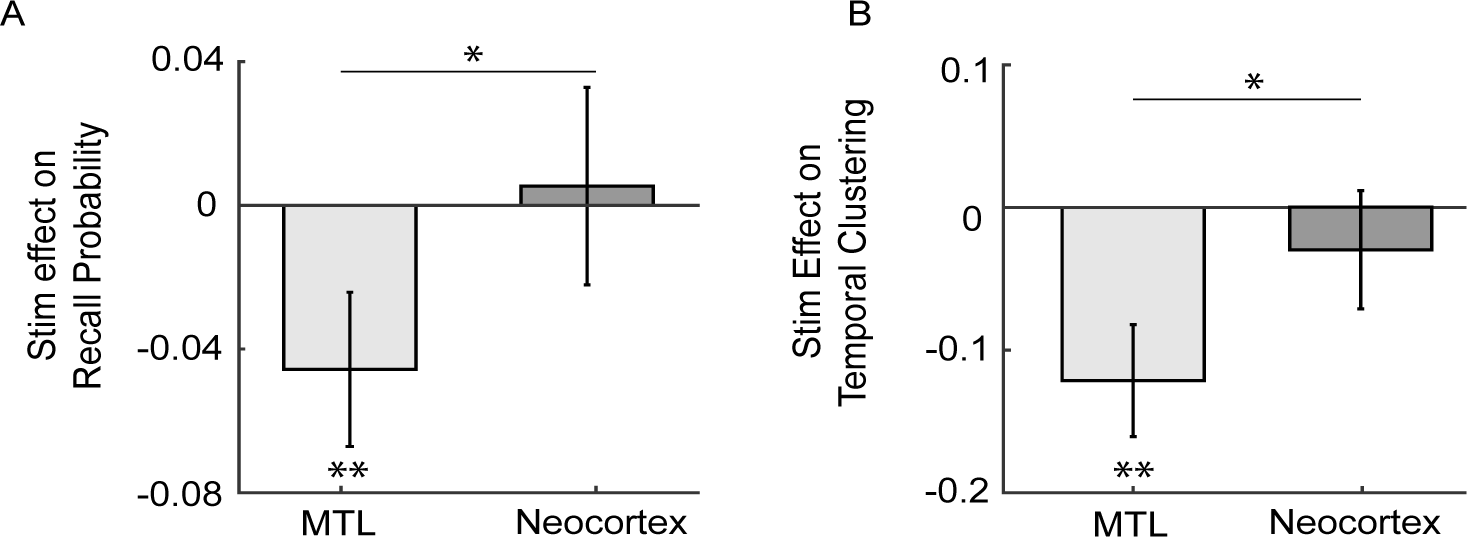
Comparison of effects of stimulation in different regions during free recall. **A.** Change in recall probability due to stimulation (stimulated lists - control lists). This measure was separately computed for stimulation that was applied in hippocampus/entorhinal cortex (labeled “MTL”) or in neocortex. **B.** Change in temporal clustering factor (stimulated lists - control lists) due to stimulation applied to MTL and neocortex regions. *: *p* < 0.05, **: *p* < 0.01

### Spatial Memory

Our spatial memory task tested subjects’ ability to remember the locations where items had been observed in a virtual reality environment. We began our analyses by examining overall task performance, as measured by our memory score (MS) measure, for subjects with stimulation electrodes in the MTL. Patients showed a range of mean memory scores, ranging from 0.51 to 0.95. Visually, the distribution of memory scores appeared to comprise more than one performance group (Fig. 6A). We determined quantitatively that splitting our subject population into two groups provided the best fit for this performance distribution using the *k*-means gap statistic (Tibshirani et al., 2001). Thus split our subjects into two performance groups—“good-performers” and “bad-performers”—using a threshold of MS=0.75.

**Figure 6:**
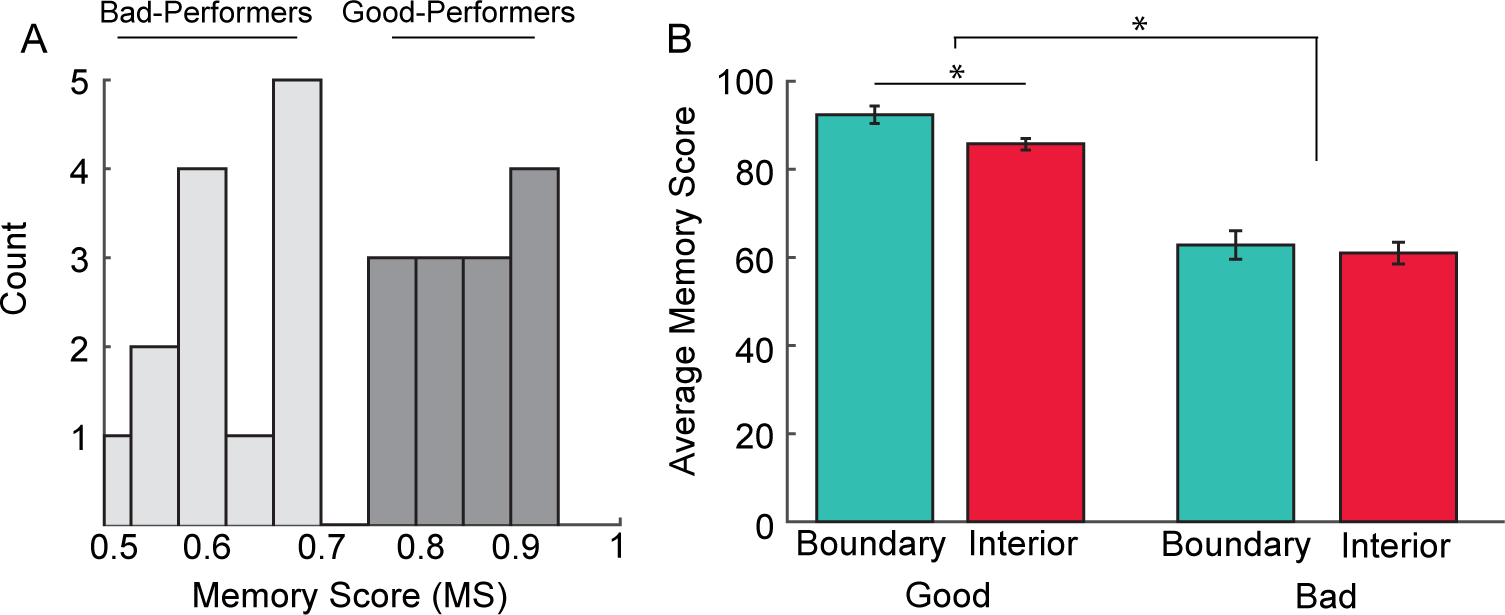
Behavioral analysis of memory performance in the spatial memory task without MTL stimulation. **A.** Histogram of memory scores across subjects with electrodes in the MTL. **B.** Average memory score for good- and bad-performing subjects, separately computed for remembered locations near and far from boundaries, then averaged across subjects. *: *p* < 0.05

We were interested in understanding the source of this performance difference. Prior work suggests that boundaries, in particular, are an important influence on spatial navigation and memory (Hartley et al., 2004; S. A. Lee, 2017; Chan et al., 2012). Furthermore, data from subjects performing this same task without stimulation showed differences in both memory performance and neural signals near boundaries (S. A. Lee et al., 2018). This body of earlier work motivated us to consider that one way the two performance groups could be distinguished is by their behavior in relation to boundaries. Thus, our subsequent analyses separately considered boundary trials and interior trials for each of the two subject performance groups.

### Assessing spatial memory strategies

We hypothesized that part of the reason that the good-performers in our task show improved performance is because they were more effective at utilizing visual information from nearby boundaries to assist with encoding object locations. We tested this by using an ANOVA to examine the effects of object location (boundary/interior, a repeated measure), subject condition group, and their interaction on memory score for trials without stimulation (Fig. 6B). Although this analysis showed that MS did not vary significantly with object location as a main factor (*F*(1, 22) = 2.16, *p* = 0.15), there was a significant interaction between subject group (i.e., good or bad) and object location (*F*(1, 22) = 4.21, *p* = 0.018). This indicated that good-performers showed significantly better memory performance near boundaries compared to bad-performers. Post-hoc tests confirmed that good-but not bad-performing subjects showed significantly greater MS for items located near boundaries (HSD-corrected post-hoc tests: good-performers, *p* = 0.047; bad-performers, *p* = 0.93).

We confirmed that this pattern was robust by analyzing a separate dataset of 69 subjects who performed the same task without stimulation (S. A. Lee et al., 2018). Here we again found that good-performers showed a significantly larger improvement in MS near boundaries than bad-performers (interaction *F*(1, 67) = 4.94, *p* = 0.028, two-way ANOVA), and that only good-performers demonstrated boundary-related performance improvements (post-hoc tests: good-performers *p* = 0.032; bad-performers *p* = 0.99). This replication of the findings from our main dataset supports the view that good-performing subjects exhibit improved memory performance when remembering locations near boundaries in this task.

The finding that one group of subjects showed increased memory performance for remembering locations near boundaries indicated to us that it was possible that these subjects varied their memory strategy for objects in different locations. We hypothesized that the subjects who showed increased memory performance for objects near boundaries employed a strategy in which they attended to visual-boundary cues during encoding and made use of those same environmental features to guide retrieval. Figure 7A presents a visualization of how this type of “visual-boundary-based” cuing could occur in this task. This technique would be more effective when subjects are near boundaries because the increased size of visual cues make them more salient. By contrast, Figure 7B depicts a “visual-boundary-agnostic” representation, in which subjects remember each location based on a global sense of their position relative to the environment. This type of representation would likely be equally useful for remembering locations both near and far from boundaries.

**Figure 7:**
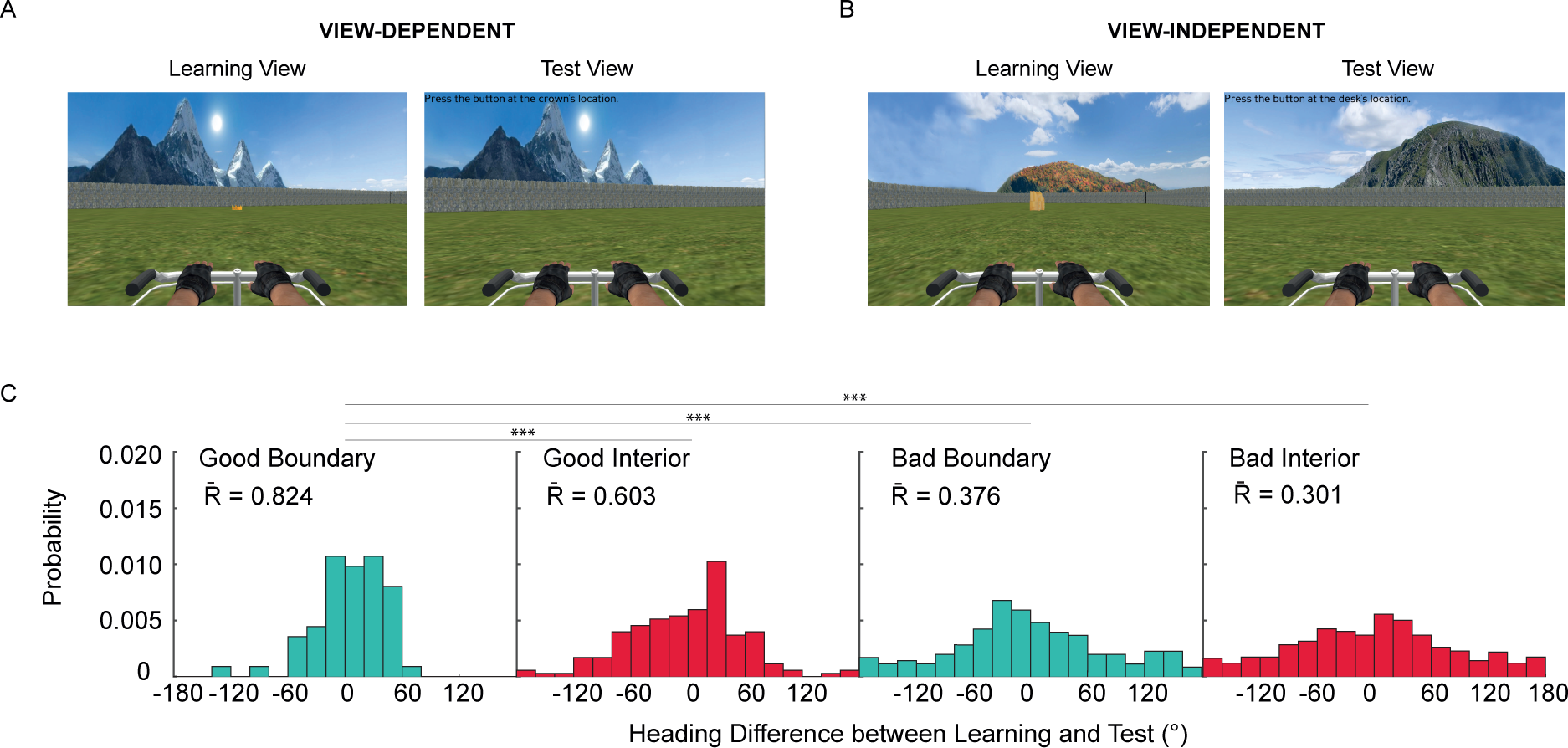
Analysis of navigational representations used by subjects independent of MTL stimulation. **A.** First-person visualization of a subject utilizing visual boundary-based cuing on a trial where they remembered a location near a boundary. **B.** Same, but for a trial where the subject does not utilize visual-boundary-based cuing on a trial where they remembered a location near the interior. **C.** Probability density functions of differences in headings between learning and test trials. Length of resultant vector (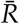) for the differences between learning and test headings for each category are also indicated. Large values imply significant clustering about 0. Heading differences were averaged across subjects in each category. ***: *p* < 0.001

Based on our finding that only good-performers showed better memory performance when remembering objects near boundaries (Fig. 6B), we hypothesized that good spatial encoding of boundary locations is likely to involve, at least in part, the use of visual-boundary-based scene representations. To test this hypothesis, we estimated the spatial representation used on each trial by measuring the difference between each subject’s heading at the end of learning and test (see *Methods*). We computed the distribution of learning–test heading differences separately for the interior and boundary regions of the environment, as well as for good- and bad-performers. If a subject had a similar heading between learning and test on a given trial, it indicates that they matched visual scene information between encoding and recall. Consistent with our predictions, good-performing subjects near boundaries showed more similar headings between learning and test compared to other memory conditions (Fig. 7C; pairwise *k* tests, all 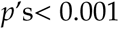). This result supports our hypothesis that good-performing subjects were more likely to utilize visual-boundary-based representations to assist with remembering objects near boundaries.

Objects located in the interior of the environment can easily be approached from any direction, whereas objects near boundaries are most often approached by driving from the center of the environment. To ensure that this aspect of the task design did not influence our results, we conducted a simulation of the mean learning–test heading differences that would be expected by chance (see *Methods*). The heading differences for good-performers near boundaries in our data were strongly clustered near zero with a mean resultant vector length 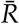 (Fig. 7C). This mean resultant vector length was well outside the range of 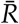 values that we observed in our randomized simulations (*p* < 0.01; simulated 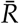 range 0.23–0.62), indicating that the learning–test heading similarities for boundary items were not artifacts of the task design.

### Effect of Stimulation

Overall, performance in the spatial memory task decreased with MTL stimulation (Jacobs et al., 2016). Given our results indicating that good-performing subjects use different memory strategies near boundaries, we assessed the effects of MTL stimulation on memory score for each subject performance group and environment area using a two-way ANOVA (Fig. 8A). This analysis showed that stimulation’s effect on MS was modulated significantly by performance group (*F*(1, 24) = 8.33, *p* < 0.01) and by object location (*F*(1, 24) = 4.55, *p* = 0.038). Thus, MTL stimulation impaired performance more in the interior of the environment and more for good-performing subjects. A post-hoc test confirmed that the memory impairment from stimulation was statistically significant for good-performing subjects in the interior of the environment (*p* = 0.010, one-sample *t* test, Bonferroni-corrected α_*crit*_ = 0.016).

**Figure 8:**
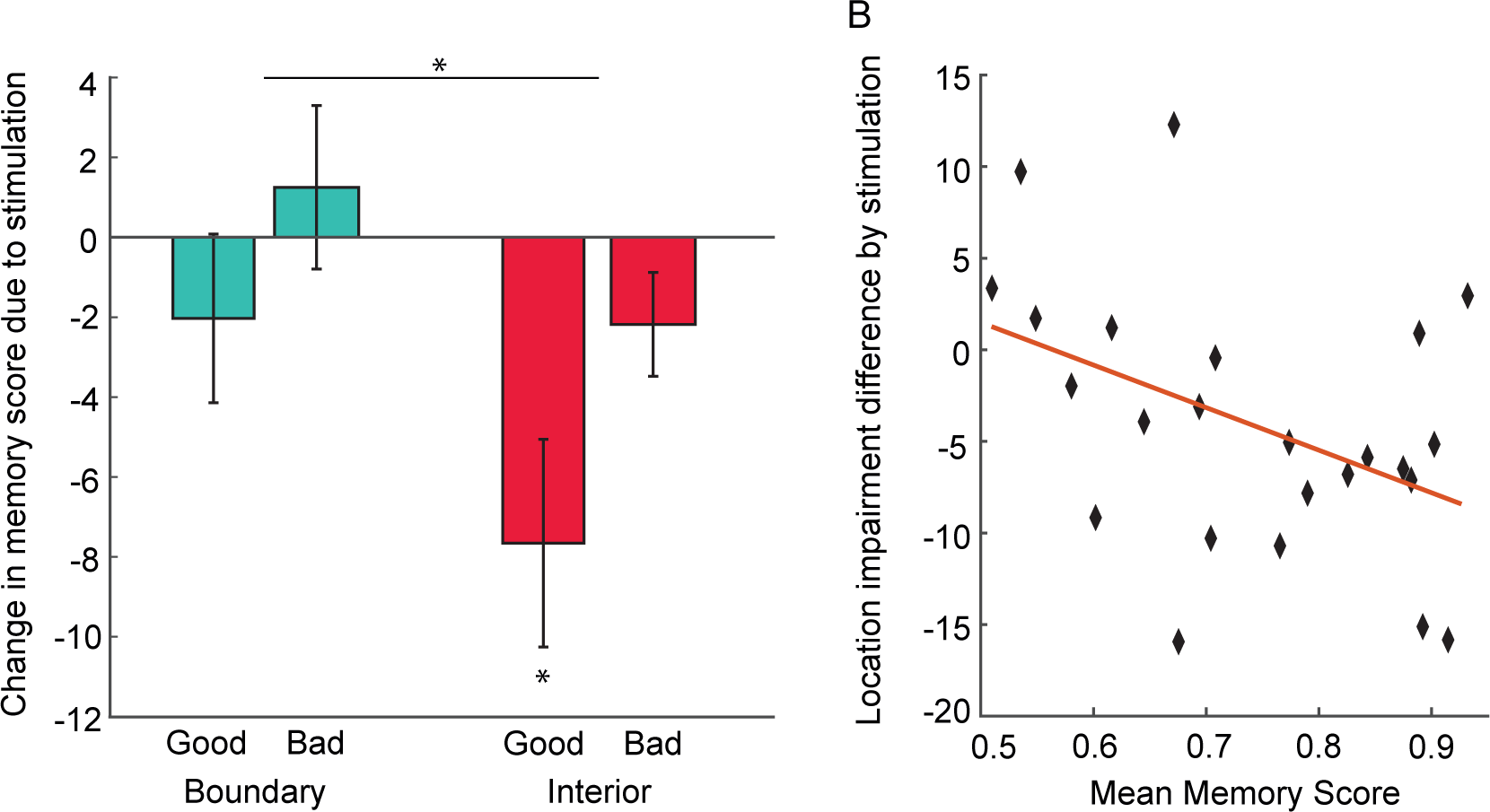
Analysis of effect of MTL stimulation on memory for objects from different locations. **A.** Difference in memory score (× 100) due to MTL stimulation for different object locations and subject types. Negative values indicate impairment from stimulation. Impairments were first averaged across trials and then across subjects. *: *p* < 0.05, †: *p* < 0.1 **B.** Scatter plot for individual subjects of the differences in memory score between stimulated and non-stimulated and between boundary and interior trials. Each point represents by how much stimulation impairs boundary trials more than interior trials. A negative value indicates increased impairment for interior trials. Line represents best-fit trend line from linear regression.

Figure 8B illustrates our pattern of results more fully by plotting the relation between each subject’s mean memory score and the differential effect of stimulation on their mean memory score for boundary versus interior trials. There is a significant negative relation between these two variables (*t*_24_ = −2.21, *p* = 0.038, one-sample *t* test), which indicates that, with increasing subject performance, stimulation caused greater memory impairment for items near the interior of the environment. These results indicate that visual-boundary-agnostic representations, which subjects likely use to remember items in the environment’s interior, were specifically impaired by MTL stimulation.

Comparing the effects of MTL stimulation on memory performance between the spatial and verbal tasks, we found a weakly positive but non-significant correlation across subjects (ρ = 0.19, *p* = 0.65). However, this analysis may be underpowered because only eight subjects contributed to this comparison.

## Discussion

By identifying the nature of the changes in memory performance that related to direct electrical brain stimulation, we have provided causal evidence to suggest that the human MTL supports the encoding of memories according to their spatial and temporal features. This bolsters our understanding of the specific cognitive processes implemented by the MTL to allow humans to maintain a framework of when and where past events occurred. By using temporally restricted methods to causally identify the means by which the MTL maintains episodic memory representations, our results provide strong evidence that is consistent with findings from previous lesion (Bohbot et al., 1998, 2004; Kolarik et al., 2017; Spiers, Maguire, & Burgess, 2001; Spiers, Burgess, Maguire, et al., 2001), imaging (Copara et al., 2014; Suthana et al., 2009; Staresina & Davachi, 2009) and intracranial EEG studies (Watrous et al., 2013; Ekstrom & Bookheimer, 2007; Miller et al., 2013; Howard et al., 2012; Foster et al., 2013).

Theoretical models have suggested that the MTL plays a role in supporting episodic memory coding with place cells and time cells (Howard & Eichenbaum, 2015). Place cells activate to encode an animal’s presence at a specific location in a given environment, representing that location in an allocentric manner relative to the overall constellation of the environment’s spatial geometry (O’Keefe & Nadel, 1978). Analogously, time cells activate at particular moments in temporally structured intervals, relative to the overall scaling of the event sequence (Eichenbaum, 2014). Based on the properties of place and time cells, it suggested that the role of the MTL, and specifically the hippocampus, in memory is to represent the overall spatiotemporal context for memory coding (Squire et al., 2004; Howard & Eichenbaum, 2015; Ekstrom & Ranganath, 2017). Our findings indicate that MTL place and time cells could be closely involved in the encoding of spatial and temporal memories (Miller et al., 2013), as the nature of the memory impairments we observed from stimulation is what would be expected if place- and time-cell representations were disrupted.

Notably, specific features of time cells correspond to aspects of our verbal memory findings. In rodents, the time cells that represent early moments in an interval each have shorter activations compared with those that represent later moments (MacDonald et al., 2011). As a result, the hippocampal population representation of temporal context evolves more rapidly at the beginning of a given interval (Howard et al., 2015). This phenomenon could relate to our finding that the disruptive effect of stimulation was stronger for primacy items. MTL stimulation may be more disruptive at the beginning of each list if the temporal context representations in this interval were more transient, or fragile, perhaps owing to the shorter-lived responses of the underlying time cells.

It is important theoretically to note that stimulation caused decreased temporal clustering (Figure 4), as evidenced by a flattened lag-CRP curve for items learned while the stimulator was active. The clustering of item recalls based on the times when they were encoded is a key element of retrieved-context models of episodic memory (Howard & Kahana, 1999); our results therefore support these models generally and indicate that the MTL has a key functional role in the neural instantiation and encoding of episodic contexts (Howard et al., 2005). Notably, stimulation in our task occurred during encoding whereas the item clustering changes we measured were during recall. Thus our findings causally show that clustering of recalls is, at least in part, a result of neural signals during encoding, perhaps due to the construction of associations from each viewed item to the current temporal context represented by time cells.

Analogous to how stimulation disrupts encoding of memory representations that may rely on MTL time cells, our data from the spatial task indicates that MTL stimulation could also disrupt the encoding of items that may rely on place and grid cells. Because the encoding of items in the interior of the environment was more strongly disrupted by stimulation, it suggests that encoding these locations was more directly supported by the MTL. Our finding that MTL stimulation does not significantly interfere with memories of objects located near boundaries does not necessarily indicate that the MTL has no role in the encoding of items near boundaries—indeed, MTL boundary cells provide input to place cells (Barry et al., 2006). Together these findings suggest that non-MTL structures can support view-based memory strategies, facilitating the encoding of certain types of memory items independently of the MTL. Consistent with this interpretation, there is evidence from fMRI studies that spatial scene recognition is mediated by both MTL and non-MTL structures, such as the striatum, parahippocampal place area, retrosplenial cortex, and occipital place area (Epstein et al., 2007; Park & Chun, 2009; Park et al., 2011; Julian et al., 2016; Doeller et al., 2008). Furthermore, it has also been directly shown that subjects with MTL damage have intact scene recognition abilities (Spiers, Burgess, Hartley, et al., 2001). These findings support our view that even when the MTL was impaired by stimulation, performance was relatively unaffected on boundary trials because extra-MTL areas were able to compensate for the deficit. Further support for this claim would arise from using a paradigm with manipulations to directly identify allocentric spatial representations, rather than our approach of inferring strategy based on mean behavior.

A potentially surprising feature of our results is that, in the spatial task, good-but not bad-performing subjects exhibited differential memory performance according to the location of an object. This emphasizes that strategy selection may be an important element of memory, for settings where multiple memory systems could potentially accomplish a task (Doll et al., 2014; Squire, 1992; Iaria et al., 2003). Because we found a correlation between subject mean performance and the tendency for stimulation to impair memory encoding far from boundaries, it suggests that good-performers were more likely to alter their spatial memory strategy depending on the object’s location relative to the boundaries. This indicates that these subjects preferentially utilized MTL-based strategies in the interior of the environment while recruiting extra-MTL brain areas for objects near boundaries to support visual-boundary-based encoding.

Our primary result is showing that the MTL is responsible for encoding the spatial and temporal structure of particular types of episodic memories. By employing causal and temporally reversible methods, we provide perhaps the strongest evidence yet for this claim. In addition to being important for our fundamental understanding of brain function and memory systems, our results have implications for guiding the future use of brain stimulation for cognitive enhancement.

Memory includes a diverse range of cognitive and neural processes. Our findings suggest that to develop brain stimulators for memory enhancement, it might be useful to tune these devices to a specific memory strategy or behavioral process. Neuromodulation in the MTL may be particularly useful for modulating memories, based on the activity of hippocampal place and time cells. Nonetheless it may still be possible that other types of neocortical stimulation could be used to modulate memory. Several types of studies have demonstrated a role for the neocortex in episodic memory, including studies with lesions (Duarte et al., 2010), direct lateral-temporal stimulation (Ezzyat et al., 2018; Kucewicz et al., 2018), and transcranial magnetic stimulation (J. X. Wang et al., 2014). Thus, probing the effects of neocortical stimulation is likely to be a focus of much future work, as it is possible that our low sample size in these regions led us to underestimate its therapeutic potential. One approach that could be useful for such enhancement is to utilize a closed-loop approach to stimulation by measuring ongoing neuronal activity in real-time and dynamically varying the nature of the stimulation that will be applied (Ezzyat et al., 2017, 2018). Given the complexity of human memory and cognition, this type of dynamic approach would be useful by allowing stimulation to vary according to instantaneous internal neural states as well as external environmental demands.

## Acknowledgements

This work was supported by the DARPA Restoring Active Memory (RAM) program (Cooperative Agreement N66001-14-2-4032) and the National Institutes of Health (MH061975). The views, opinions, and/or findings expressed are those of the authors and should not be interpreted as representing the official views or policies of the Department of Defense or the U.S. Government. We thank Blackrock Microsystems for providing neural recording and stimulation equipment. We thank Michael Kahana and Daniel Rizzuto for assisting with data collection and for scientific discussions.

## References

Avants, B. B., Epstein, C. L., Grossman, M., & Gee, J. C. (2008). Symmetric diffeomorphic image registration with cross-correlation: evaluating automated labeling of elderly and neurodegenerative brain. Medical Image Analysis, 12(1), 26–41.

Barry, C., Lever, C., Hayman, R., Hartley, T., Burton, S., O’Keefe, J.,…Burgess, N. (2006). The boundary vector cell model of place cell firing and spatial memory. Rev Neurosci, 17(1-2), 71–97.

Berens, P. (2009). Circstat: A matlab toolbox for circular statistics. Journal of Statistical Software, 31(10).

Bohbot, V., Iaria, G., & Petrides, M. (2004). Hippocampal function and spatial memory: evidence from functional neuroimaging in healthy participants and performance of patients with medial temporal lobe resections. Neuropsychology, 18(3), 418–25.

Bohbot, V., Kalina, M., Stepankova, K., Spackova, N., Petrides, M., & Nadel, L. (1998). Spatial memory deficits in patients with lesions to the right hippocampus and to the right parahippocampal cortex. Neuropsychologia, 36(11), 1217–38.

Chan, E., Baumann, O., Bellgrove, A., Mark, & Mattingley, B., Jason. (2012). From objects to landmarks: the function of visual location information in spatial navigation. Frontiers in Psychology, 3(304).

Cohen, N. J., & Eichenbaum, H. (1993). Memory, amnesia, and the hippocampal system. Cambridge, MA: MIT.

Copara, M. S., Hassan, A., Kyle, C. T., Libby, L., Ranganath, C., & Ekstrom, A. D. (2014). Complementary roles of human hippocampal subregions during retrieval of spatiotemporal context. Journal of Neuroscience, 34(20).

Craik, F. I. M. (1970). The fate of primary memory items in free recall. Journal of Verbal Learning and Verbal Behavior, 9, 658–664.

Darley, C. F., & Murdock, B. B. (1971). Effects of prior free recall testing on final recall and recognition. Journal of Experimental Psychology, 91, 66–73.

Doeller, C. F., King, J. A., & Burgess, N. (2008). Parallel striatal and hippocampal systems for landmarks and boundaries in spatial memory. Proceedings of the National Academy of Sciences, USA, 105(15), 5915–5920.

Doll, B., Shohamy, D., & Daw, N. (2014). Multiple memory systems as substrates for multiple decision systems. Neurobiology of learning and memory.

Douglas, D., Thavabalasingham, S., Chorghay, Z., O’Neil, E. B., Barense, M. D., & Lee, A. C. H. (2016). Perception of impossible scenes reveals differential hippocampal and parahippocampal place area contributions to spatial coherency. Hippcampus, 27(1).

Duarte, A., Henson, R., Knight, R. T., Emery, T., & Graham, K. S. (2010). The orbitofrontal cortex is necessary for temporal context memory. Journal of Cognitive Neuroscience, 22(8), 1819–1831.

Eichenbaum, H. (2004, Sep). Hippocampus: cognitive processes and neural representations that underlie declarative memory. Neuron, 44(1), 109–120. Retrieved from http://dx.doi.org/10.1016/j.neuron.2004.08.028 doi: 10.1016/j.neuron.2004.08.028

Eichenbaum, H. (2014). Time cells in the hippocampus: a new dimension for mapping memories. Nature Reviews Neuroscience, 15(11), 732–744.

Ekstrom, A. D., & Bookheimer, S. Y. (2007). Spatial and temporal episodic memory retrieval recruit dissociable functional networks in the human brain. Learning & Memory.

Ekstrom, A. D., Copara, M. S., Isham, E. A., Wang, W., & Yonelinas, A. P. (2011). Dissociable networks involved in spatial and temporal order source retrieval. NeuroImage, 56(3), 1803–1813.

Ekstrom, A. D., & Ranganath, C. (2017). Space, time and episodic memory: the hippocampus is all over the cognitive map. Hippocampus.

Epstein, R. A., Parker, W., & Feiler, A. (2007). Where am I now? Distinct roles for parahippocampal and retrosplenial cortices in place recognition. Journal of Neuroscience, 27(23), 6141.

Ezzyat, Y., Kragel, J. E., Burke, J. F., Levy, D. F., Lyalenko, A., Wanda, P.,…Kahana, M. J. (2017). Direct brain stimulation modulates encoding states and memory performance in humans. Current Biology, 27, 1–8.

Ezzyat, Y., Wanda, P., Levy, D., Kadel, A., Aka, A., Pedisich, I.,…Kahana, M. (2018). Closed-loop stimulation of temporal cortex rescues functional networks and improves memory. Nature Communications, 9(365). doi: 10.1038/s41467-017-02753-0

Fischler, I., Rundus, D., & Atkinson, R. C. (1970). Effects of overt rehearsal procedures on free recall. Psychonomic Science, 19(4), 249–250.

Foster, B. L., Kaveh, A., Dastjerdi, M., Miller, K. J., & Parvizi, J. (2013). Human retrosplenial cortex displays transient theta phase locking with medial temporal cortex prior to activation during autobiographical memory retrieval. The Journal of Neuroscience, 33(25), 10439–10446.

Friston, K. J., Glaser, D., Henson, R. N. A., Kiebel, S., Phillips, C., & Ashburner, J. (2002). Classical and bayesian inference in neuroimaging: applications. NeuroImage, 16.

Guderian, S., Dzieciol, A. M., Gadian, D. G., Jentschke, S., Doeller, C. F., Burgess, N.,…Vargha-Khadem, F. (2015). Hippocampal volume reduction in humans predicts impaired allocentric spatial memory in virtual-reality navigation. Journal of Neuroscience, 35(42).

Hartley, T., Trinkler, I., & Burgess, N. (2004). Geometric determinants of human spatial memory. Cognition, 94, 39–75.

Henson, R. (2005). A mini-review of fmri studies of human medial temporal lobe activity associated with recognition memory. The Quarterly Journal of Experimental Psychology, 58(3-4).

Howard, M. W., & Eichenbaum, H. (2015). Time and space in the hippocampus. Brain Research.

Howard, M. W., Fotedar, M. S., Datey, A. V., & Hasselmo, M. E. (2005). The temporal context model in spatial navigation and relational learning: Toward a common explanation of medial temporal lobe function across domains. Psychological Review, 112(1), 75–116.

Howard, M. W., & Kahana, M. J. (1999). Contextual variability and serial position effects in free recall. Journal of Experimental Psychology: Learning, Memory, and Cognition, 25(4), 923–941. doi: 10.1037/0278-7393.25.4.923

Howard, M. W., & Kahana, M. J. (2002). A distributed representation of temporal context. Journal of Mathematical Psychology, 46(3), 269–99.

Howard, M. W., Shankar, K. H., Aue, W. R., & Criss, A. H. (2015). A distributed representation of internal time. Psychological Review, 122(1), 24–53. doi: 10.1037/a0037840

Howard, M. W., Viskontas, I. V., Shankar, K. H., & Fried, I. (2012). Ensembles of human MTL neurons "jump back in time" in response to a repeated stimulus. Hippocampus, 22, 1833–1847.

Iaria, G., Petrides, M., Dagher, A., Pike, B., & Bohbot, V. (2003). Cognitive strategies dependent on the Hippocampus and Caudate Nucleus in human navigation: Variability and change with practice. Journal of Neuroscience, 23(13), 5945–5952.

Jacobs, J., Miller, J., Lee, S. A., Coffey, T., Watrous, A. J., Sperling, M. R.,…Rizzuto, D. S. (2016, December). Direct electrical stimulation of the human entorhinal region and hippocampus impairs memory. Neuron, 92(5), 1–8.

Julian, J. B., Hamilton, R. H., & Epstein, R. A. (2016). The occipital place area is causally involved in representing environmental boundaries during navigation. Current Biology, 26, 1104–1109.

Kahana, M. J. (1996). Associative retrieval processes in free recall. Memory & Cognition, 24(1), 103–109.

Kahana, M. J. (2012). Foundations of human memory. New York, NY: Oxford University Press.

Klomjai, W., Katz, R., & Lackmy-Valee, A. (2015). Basic principles of transcranial magnetic stimulation (tms) and repetitive tms (rtms). Annals of Physical and Rehabilitation Medicine, 58(4), 208–213.

Knotkova, H., & Rasche, D. (2014). Textbook of neuromodulation: Principles, methods, and clinical applications. Springer Science & Business Media.

Kolarik, B. S., Baer, T., Shahlaie, K., Yonelinas, A. P., & Ekstrom, A. D. (2017). Close but no cigar: spatial precision deficits following medial temporal lobe lesions provide novel insight into theoretical models of navigation and memory. Hippocampus.

Kucewicz, M. T., Berry, B., Miller, L., Khadjevand, F., Ezzyat, Y., Stein, J.,…Worrell, G. (2018). Evidence for verbal memory enhancement with electrical brain stimulation in the lateral temporal cortex. Brain. doi: 10.1093/brain/awx373

Lee, S. A. (2017). The boundary-based view of spatial cognition: a synthesis. Current Opinion in Behavioral Sciences, 16, 58–65.

Lee, S. A., Miller, J. F., Watrous, A. J., Sperling, M., Sharan, A., Worrell, G. A.,…others (2018). Electrophysiological signatures of spatial boundaries in the human subiculum. Journal of Neuroscience, 218040.

Lee, S. H., & Dan, Y. (2012). Neuromodulation of brain states. Neuron, 76(1), 209–222.

MacDonald, C., Lepage, K., Eden, U., & Eichenbaum, H. (2011). Hippocampal "time cells" bridge the gap in memory for discontiguous events. Neuron, 71(4), 737–749.

Maguire, E. A., Intraub, H., & Mullally, S. L. (2015). Scenes, spaces, and memory traces. what does the hippocampus do? The Neuroscientist, 22(5).

Manning, J. R., Polyn, S. M., Baltuch, G., Litt, B., & Kahana, M. J. (2011). Oscillatory patterns in temporal lobe reveal context reinstatement during memory search. Proceedings of the National Academy of Sciences, USA, 108(31), 12893–12897. doi: 10.1073/pnas.1015174108

Miller, J. F., Neufang, M., Solway, A., Brandt, A., Trippel, M., Mader, I.,…Schulze-Bonhage, A. (2013). Neural activity in human hippocampal formation reveals the spatial context of retrieved memories. Science, 342(6162), 1111–1114.

Morris, R. (1984). Developments of a water-maze procedure for studying spatial learning in the rat. Journal of Neuroscience Methods, 11(1), 47–60.

Murdock, B. B. (1962). The serial position effect of free recall. Journal of Experimental Psychology, 64, 482–488. doi: 10.1037/h0045106

O’Keefe, J., & Dostrovsky, J. (1971). The hippocampus as a spatial map: Preliminary evidence from unit activity in the freely-moving rat. Brain Research, 34, 171–175.

O’Keefe, J., & Nadel, L. (1978). The hippocampus as a cognitive map. New York: Oxford University Press.

Park, S., Brady, T., Greene, M., & Oliva, A. (2011). Disentangling scene content from spatial boundary: Complementary roles for the parahippocampal place area and lateral occipital complex in representing real-world scenes. Journal of Neuroscience, 31, 1333–1340.

Park, S., & Chun, M. M. (2009). Different roles of the parahippocampal place area (ppa) and retrosplenial cortex (rsc) in panoramic scene perception. NeuroImage, 47(4), 1747–1756.

Polyn, S. M., Norman, K. A., & Kahana, M. J. (2009a). A context maintenance and retrieval model of organizational processes in free recall. Psychological Review, 116, 129–156. doi: 10.1037/a0014420

Polyn, S. M., Norman, K. A., & Kahana, M. J. (2009b). Task context and organization in free recall. Neuropsychologia, 47, 2158–2163. doi: 10.1016/j.neuropsychologia.2009.02.013

Ramsey, J., S.J., H., C., H., Halchenko, Y. O., Poldrack, R. A., & Glymour, C. (2010). Six problems for causal inference from fmri. NeuroImage, 49(2).

Rorden, C., & Karnath, H.-O. (2004). Using human brain lesions to infer function: a relic from a past era in the fmri age? Nature Reviews Neuroscience.

Scoville, W. B., & Milner, B. (1957). Loss of recent memory after bilateral hippocampal lesions. Journal of Neurology, Neurosurgreery, and Psychiatry, 20, 11–21.

Sederberg, P. B., Gauthier, L. V., Terushkin, V., Miller, J. F., Barnathan, J. A., & Kahana, M. J. (2006). Oscillatory correlates of the primacy effect in episodic memory. NeuroImage, 32(3), 1422–1431. doi: 10.1016/j.neuroimage.2006.04.223

Sederberg, P. B., Kahana, M. J., Howard, M. W., Donner, E. J., & Madsen, J. R. (2003). Theta and gamma oscillations during encoding predict subsequent recall. Journal of Neuroscience, 23(34), 10809–10814.

Serruya, M. D., Sederberg, P. B., & Kahana, M. J. (2014). Power shifts track serial position and modulate encoding in human episodic memory. Cerebral Cortex, 24, 403–413. doi: 10.1093/cercor/bhs318

Spiers, H. J., Burgess, N., Hartley, T., Vargha-Khadem, F., & O’Keefe, J. (2001). Bilateral hippocampal pathology impairs topographical and episodic memory but not visual pattern matching. Hippcampus, 11(6).

Spiers, H. J., Burgess, N., Maguire, E. A., Baxendale, S. A., Hartley, T., Thompson, P. J., & O’Keefe, J. (2001). Unilateral temporal lobectomy patients show lateralized topographical and episodic memory deficits in a virtual town. Brain, 124(Pt 12), 2476–89.

Spiers, H. J., Maguire, E. A., & Burgess, N. (2001). Hippocampal amnesia. Neurocase, 7, 357–382.

Squire, L. (1992). Memory and the hippocampus: A synthesis from findings with rats, monkeys, and humans. Psychological Review, 99, 195–231.

Squire, L., Stark, C., & Clark, R. (2004). The medial temporal lobe. Annual Review of Neuroscience, 27, 279–306.

Staresina, B. P., & Davachi, L. (2009). Mind the gap: binding experiences across space and time in the human hippocampus. Neuron, 63(2), 267–276.

Suthana, N., Ekstrom, A. D., Moshirvaziri, S., Knowlton, B. J., & Bookheimer, S. Y. (2009). Human hippocampal ca1 involvement during allocentric encoding of spatial information. Journal of Neuroscience.

Suthana, N., & Fried, I. (2014). Deep brain stimulation for enhancement of learning and memory. Neuroimage, 85, 996–1002.

Suthana, N., Haneef, Z., Stern, J., Mukamel, R., Behnke, E., Knowlton, B., & Fried, I. (2012). Memory enhancement and deep-brain stimulation of the entorhinal area. The New England Journal of Medicine, 366, 502–10.

Tibshirani, R. J., Walther, G., & Hastie, T. (2001). Estimating the number of clusters in a dataset via the gap statistic. Journal of the Royal Statistical Society: Series B (Statistical Methodology), 63(2), 411–423.

Wang, H., Suh, J. W., Das, S. R., Pluta, J. B., Craige, C., Yushkevich, P., et al. (2013). Multi-atlas segmentation with joint label fusion. Pattern Analysis and Machine Intelligence, IEEE Transactions on, 35(3), 611–623.

Wang, J. X., Rogers, L. M., Gross, E. Z., Ryals, A. J., Dokucu, M. E., Brandstatt, K. L.,…Voss, J. L. (2014, August). Targeted enhancement of cortical-hippocampal brain networks and associative memory. Science, 345(6200), 1054–1057.

Watrous, A. J., Tandon, N., Conner, C. R., Pieters, T., & Ekstrom, A. D. (2013). Frequency-specific network connectivity increases underlie accurate spatiotemporal memory retrieval. Nature Neuroscience, 16(3), 349–356.

Yushkevich, P. A., Pluta, J. B., Wang, H., Xie, L., Ding, S.-L., Gertje, E. C.,…Wolk, D. A. (2015). Automated volumetry and regional thickness analysis of hippocampal subfields and medial temporal cortical structures in mild cognitive impairment. Human Brain Mapping, 36(1), 258–287.

